# Structure of *Escherichia coli* DNA gyrase with chirally wrapped DNA supports ratchet-and-pawl mechanism for an ATP-powered supercoiling motor

**DOI:** 10.1101/2024.04.12.589215

**Authors:** Elizabeth Michalczyk, Marta Pabiś, Jonathan Heddle, Dmitry Ghilarov

## Abstract

Gyrase is essential for replication and transcription in bacteria, and as such is an important target for antibiotics including fluoroquinolones. Gyrase is a molecular machine that channels the energy of ATP hydrolysis into negative supercoiling of DNA. The mechanism proposed more than 40 years ago involves stabilising and inverting a chiral DNA loop; however, the molecular basis for this is poorly understood, as the loop was never directly observed. We present high-resolution cryoEM structures of the *Escherichia coli* gyrase - 217 bp DNA holocomplex, and of the moxifloxacin-bound gyrase complex with cleaved 217 bp DNA. Each structure constrains an intact figure-of-eight positively supercoiled DNA loop, poised for strand passage. The loop is stabilised by a GyrA β-pinwheel domain which how we here show has a structure of a flat disc, and functions akin to a mini-nucleosome. Our data implies that during catalysis the ATPase domains of the enzyme undergo a large movement to push the transported DNA segment through the break in DNA. By comparing the catalytic site between native drug-free and moxifloxacin-bound gyrase structures that both contain a single metal ion we demonstrate that the enzyme is observed in a native pre-catalytic state. Finally, we propose a ‘ratchet and pawl’ mechanism for energy coupling in gyrase. These unexpected findings call for re-evaluation of existing data and offer a framework for further experiments designed to dissect the details of how gyrase molecular motor converts chemical energy into mechanical tension.

## Main text

Molecular machines consume energy, typically in the form of nucleotide triphosphates, to overcome thermal fluctuations and produce unidirectional motion. A few well-studied examples include kinesin, myosin and F1-ATPase; however, how exactly localised energy consumption in the form of nucleotide binding and release results in nm-scale directional movements remains a fundamental question^1^. Understanding of the organisational principles of molecular machines is important for the manipulation of their activities and design of artificial nature-inspired nanoscale devices.

DNA gyrase is a bacterial type II topoisomerase belonging to the gyrase-Hsp90-kinase-MutL (GHKL) ATPase family: members of this group (topoisomerases, DNA repair proteins, heat shock proteins, and, recently, prokaryotic and eukaryotic immunity proteins) use ATP to trigger dimerization and transition through distinct conformational steps transducing energy into mechanistic outcomes^2–4^. Gyrase is essential in bacteria for both removing positively supercoiled DNA in front of the progressing RNA polymerase and introducing negative supercoiling required for chromosomal homeostasis. It directly and indirectly affects virtually all genomic transactions in the cell^5^. As such, gyrase is also a successful target for antibiotics with fluroquinolones being the most clinically important group.

*E. coli* gyrase is a heterotetramer formed of two GyrA and two GyrB subunits (A_2_B_2_). The GyrA subunit consists of a N-terminal winged-helix domain (WHD) and Tower domain, a long coiled-coil domain, and a C-terminal beta-pinwheel domain. The GyrB subunit comprises an N-terminal GHKL domain, a transducer hinge, and a topoisomerase-primase (Toprim) domain, with a species-specific insertion (see **Figure 1a**). GyrA subunits dimerise to form two interfaces called ‘gates’: the DNA-gate, and the C-gate. The Toprim domains of GyrB associate with GyrA to form a DNA-binding interface, while the GHKL domains are thought to be highly flexible and power DNA movements through the enzyme. While multiple crystal structures are available for isolated GHKLs and ‘core’ (GyrB: Toprim and insertion; GyrA: WHD, Tower and coiled-coil) domains of the enzyme, there are only a handful of structural studies of full-length gyrase. Two existing cryoEM structures of *E. coli* gyrase in complex with inhibitors^6,7^ display the dimerised GHKLs above the DNA-gate forming a third (ATPase) ‘gate’, while the crystal structure of DNA-free *Mycobacterium tuberculosis* gyrase revealed a backwards-bent conformation of GHKL domains which was proposed to be an energy-saving resting state stabilised by a species-specific (for *Corynebacteria*) insertion in the Toprim domain^8^.

**Figure 1.**
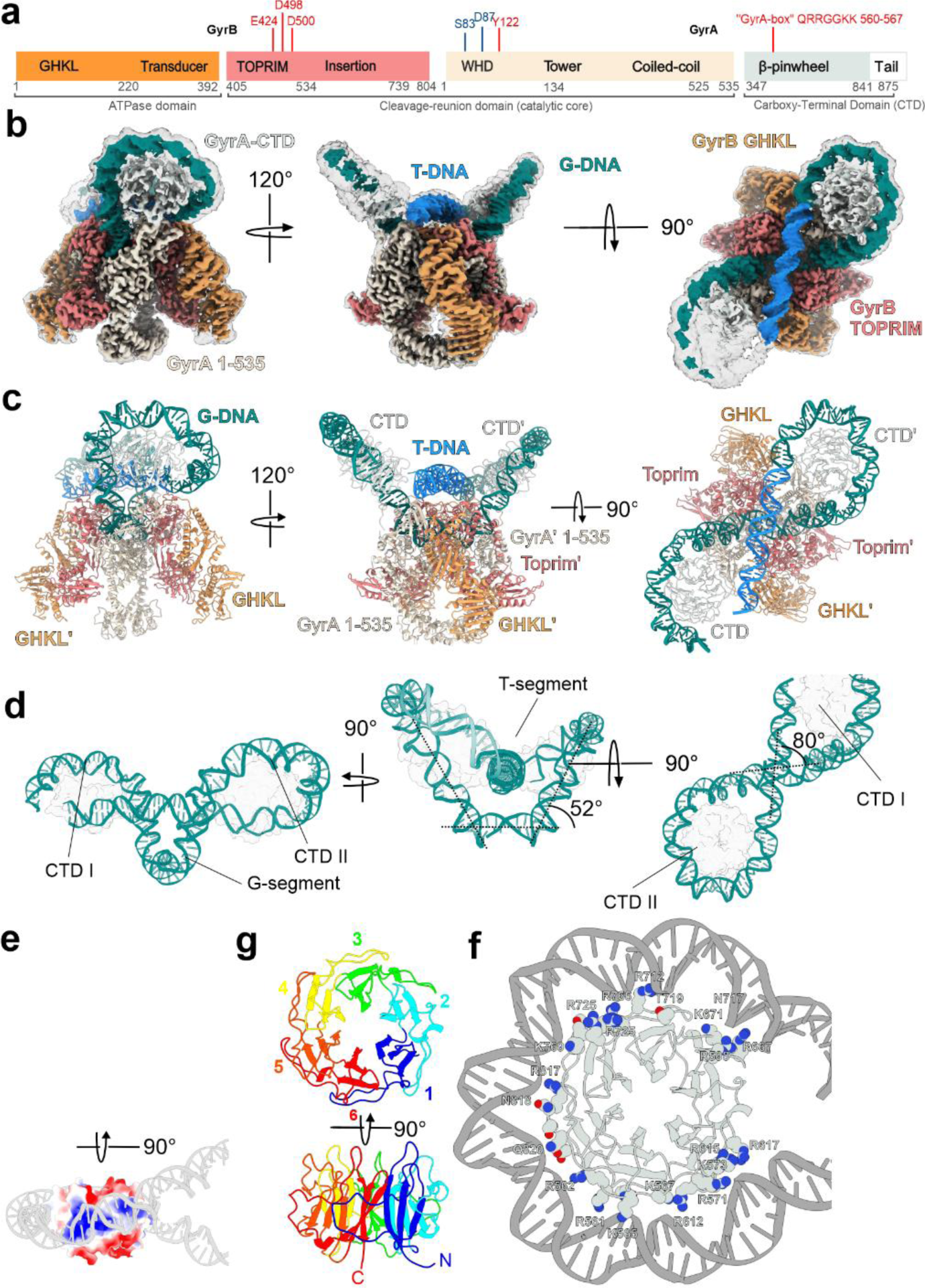
Overall architecture of chirally-wrapped *E. coli* gyrase complex & structure of a positively supercoiled DNA loop. **a**, Domain structure of GyrB (*left*) and GyrA (*right*) subunits. Key amino-acid residues important for catalysis and metal binding (red) and fluroquinolone resistance (blue) are indicated. The same color scheme is used throughout the manuscript: GHKL – orange, TOPRIM – coral, GyrA core region – beige, GyrA CTD & tail – mint white. **b**, Different views of the cryoEM density map for the **Gyr-Mu217** complex presented at two contour levels (10σ and 5σ). The sharpened 10σ map is colored according to the color scheme above but the G-segment DNA is colored teal and the T-segment DNA light blue. **c**, Corresponding views of the atomic model of complete **Gyr-Mu217** complex; protein and DNA are shown in cartoon representation. **d**, Modelled DNA loop shown in isolation with both CTDs shown as transparent contours. Angles between DNA segments are indicated. **e**, Surface representation of an isolated CTD colored by Coulombic potential (blue – positive, red – negative, measured by ChimeraX) with the Mu217 right arm wrapped around and shown as a cartoon representation. **g**, An atomic model of a CTD with blades 1 to 6 colored blue, cyan, green, yellow, orange and red respectively. **f**, A cartoon representation of the CTD. Residues within 4 Å of DNA are shown as VDW spheres. Note that GyrA box residues of each blade are interacting with DNA minor groove.

Gyrase is widely believed to operate by a tightly coordinated strand-passage node inversion mechanism^9,10^. It is conceived that during its catalytic cycle, gyrase binds a double-stranded (ds) segment of DNA called the gate-, or G-segment across the DNA gate interface and chirally wraps ∼140 bp of flanking DNA around the GyrA CTDs. Dimerisation of the ATP-controlled clamp is proposed to capture the proximal dsDNA segment called the T (for transported) segment in the cavity between the GHKLs. Subsequent G-DNA cleavage is controlled by metal binding to the GyrB Toprim domain and involves transfer of a 5’ phosphate of each DNA chain to the corresponding tyrosine (Y122) residue within the WHD domain of GyrA. It allows the proposed opening of the enzyme, leading to the movement of the trapped T-segment through the break, inverting the node and introducing two negative supercoils. DNA can then exit the enzyme via the C-gate. The catalytic reaction of the *E. coli* enzyme was shown to require additional regulatory elements: the unstructured acidic C-terminus of GyrA (‘acidic tail’)^11,12^, controlling DNA wrap, and the large insertion in the Toprim domain of the enzyme^13^.

The strand-passage model is supported by experiments showing that interface cross-linking prevents catalysis^14,15^; however, alternative modes of operation were proposed based on the fact that a mutant enzyme complex with only one catalytic tyrosine remains catalytically competent^16^. In addition, the T-segment DNA has never been visualised in any type II topoisomerase structure to date.

Here we present the first high-resolution (2.3-Å) cryoEM structure of an intact complete *E. coli* gyrase holoenzyme in the chirally wrapped state bound to 217 bp linear DNA fragment, and the parallel structure in complex with the fluoroquinolone moxifloxacin (MFX). We describe protein-DNA interactions controlling the wrapping of DNA around the CTDs that present the T-segment DNA above and perpendicular to the G-segment. Unexpectedly, the structure shows both GHKL domains folded down towards the sides of the enzyme, a conformation stabilised by multiple interactions with the Toprim insert, indicating that the nucleotide binding induces a large conformation shift. GHKL dimerization, as observed in the previous AMP-PNP bound structures, is incompatible with the position of the T-DNA, and can only happen *after* strand passage took place. that has a potential to push the T-DNA through the break, powering strand-passage. By examining the catalytic site in both subunits of the drug-free and MFX-bound complexes we demonstrate that the drug-free structure is in a pre-cleaved state different from the ones previously observed.

### In nucleotide-free conditions, *E. coli* DNA gyrase stabilises a positively supercoiled DNA loop

In order to better understand the sequence of events enabling gyrase to function as a molecular motor, we have conducted a systematic cryoEM investigation of the enzyme in different stages of its catalytic cycle, using our established 217 bp dsDNA substrate from the coliphage Mu strong gyrase site^17^ that is sufficient to support effective strand passage^7,18^. In the existing structure with a DNA-intercalating molecule albicidin, the DNA-binding CTD domains partially wrap DNA and project the ends of the linear fragment at angles that are not compatible with supercoiling directionality (the projected DNA crossing occurs below the DNA-gate axis, resulting in a negatively supercoiled DNA loop). While the previous cryoEM study^6^ refers to this conformation as ‘chirally wrapped’, we notice that in fact it is almost symmetrical and consistent with the reported Ω state that bends DNA without T-segment capture^19,20^. According to the available single-molecule and ensemble data, we have anticipated that while AMP-PNP is known to release the DNA wrap, in the absence of nucleotide and low force conditions, a chirally wrapped α state predominates ^19^. Following this, we have collected a targeted dataset **Gyr-Mu217**, processing of which resulted in a 2.3 Å consensus map displaying striking differences to the published gyrase structures, which are visible starting from the 2D class-averages level (**Figure 1b**, **Supplementary Figure 1**, **Supplementary Table 1**). Most notably, a linear DNA fragment was found to be fully wrapped around both CTDs forming a figure of eight-like contiguous positively supercoiled DNA loop that dominated the structure. To allow this, the CTD domains of the enzyme moved upwards to form a larger angle with the G-DNA plane (**Figure 1c,d**). The loop was fully modelled using the available DNA sequence, and the fact that the unique MuSGS properties position the enzyme on DNA uniformly in a defined register and orientation, resulting in an SGS ‘right arm’ chirally wrapped around one of the CTDs (CTD II) to present a T-segment for the strand passage as previously established in footprinting experiments^21^. The observed size of the loop is 156 bp, which is very close to the experimentally proposed values of the minimal length of DNA known to present the T-segment and thus stimulate strand passage^18,22^. The wrap around the opposite CTD (CTD I) was incomplete, with the remaining nucleotides of the left arm pointing away from the enzyme. The T-segment is positioned ∼2 nm above the G-segment and is almost perpendicular to it (80° angle, **Figure 1d**). Positively charged residues on both GyrA CTDs and Tower domains and GyrB Toprim domains delineate a ‘guiding path’ that can only accommodate one T-segment at a time (**Supplementary Figure 2a**). This guiding path includes a band of positive charge spiralling along the GyrA CTD residues that act as DNA-binding pulleys (**Figure 1e**). Previous X-ray crystallography analysis established that the isolated *E. coli* gyrase CTD is an incomplete β-pinwheel domain that forms a spiral structure; this spiral was proposed to be crucial for chiral loop stabilisation ^23^. In a previous cryoEM study with an incompletely wrapped DNA^6^, low resolution prevented accurate modelling of the CTD structure, as blade I was not accurately predicted by Phyre2 or Alphafold 2. In our work, we used focussed classification and refinement approach (see **Methods**) to accurately reconstruct and refined the CTD (2.9 Å) which is found to have a perfect β-pinwheel fold^24^ for all 6 blades (**Figure 1f; Supplementary Figure 2b**), in full alignment with the originally published crystal structure of *Borrelia burgdorferi* CTD. Each of the blades donates a loop that wraps around the (n-1) blade. These loops contain positively charged residues forming the so-called GyrA-box motif^25^; this motif has a different degree of conservation in each blade. GyrA-box residues are interacting with the minor groove of DNA to stabilise 5 sharp bends to convey an overall ∼260° bend, therefore each GyrA-box contributes ∼45-60° of bend (see **Supplementary Figure 2c**). GyrA-boxes of blades 1(QRRGGKK) and 2 (TRGARGR) contain the largest number of positively charged residues concomitant to their role in interacting with the proximal (CTDII) or distal (CTDI) ends of the T-segment, and maintaining it in the strand passage position (**Figure 1g**). The GyrA-box of blade 1 is a hallmark feature of all gyrases and is absolutely required for supercoiling and for the T-segment presentation; thus, we conclude that the observed supercoiled loop is the key pre-catalytic intermediate characteristic of all gyrases. Mu phage SGS is critical for the phage DNA replication cycle and is known to bind DNA strongly and support faster supercoiling. Our model demonstrates that as was hypothesized previously^21^, right arm of Mu SGS displays AT repeats located in the minor groove facing the protein surface, while GC repeats face outwards (**Supplementary Figure 2d**). The same sequence preferences are shown by the nucleosomes^26^. Strikingly, the repeated AT/GC pattern of gyrase binding can be observed on the genome-level by analysing gyrase binding site consensus^27^. Thus, the propensity of DNA to wrap around the CTDs controls gyrase location on DNA. The similarity with the nucleosome is further exacerbated by the conserved acidic tail of CTD (not observed in our structure) which was shown to be a critical element of the *E. coli* gyrase supercoiling mechanism^11^. This raises intriguing possibilities of a post-translational modification control of gyrase activity.

### GHKL domains undergo a large conformational change during catalysis

Another central feature of the nucleotide-free structure (**Figure 1c**) is the conformation of the GHKL domains, that are ‘folded down’ such that each GHKL interacts with the Toprim insertion domain of the same GyrB subunit, reminiscent of the X-ray crystallographic structure of *M. tuberculosis* gyrase^8^ or *Streptococcus pneumoniae* topoisomerase IV structure ^28^. However, both of these structures where superimposed with the Gyr-Mu217 demonstrate that T-segment position sterically clashes with the conformation of GHKL in these enzymes (**Supplementary Figure 3**). Therefore, it is not clear if in those cases the ‘folded’ conformation directly precedes supercoiling, or it used for enzyme storage as suggested^8^. In contrast, it seems that the ‘folded’ configuration of GHKLs is a native feature of at least some gyrases which evolved to control coupling of ATP binding with supercoiling. Superposition of the chirally-wrapped structure with the ADPNP-bound structure (PDB:6RKW ^6^) shows a dramatic almost 180° rotation and 12 nm shift in the position of GHKLs upon nucleotide binding (**Figure 2a**,). Importantly, the dimeric ATPase ‘clamp’ which was for decades hypothesized to contain T-DNA, sterically clashes with the T-segment position. At the same time, a single GHKL subunit if taken separately is placed comfortably on top of the T-DNA. Interestingly, this superposition places wrapped DNA in contact with the positively charged outer surface of GHKL (**Supplementary Figure 4a**). It also shows similarity to the observed interactions between GHKLs and antibiotic resistance pentapeptide repeat proteins QnrB1 and MfpA^29,30^ that were shown to mimic T-segment DNA (**Supplementary Figure 4bcd**). Nevertheless, movement to this position will require each subunit to rotate and cross the path of the T-segment, and therefore would not possible before DNA-gate opening and T-segment moving beyond the G-segment plane.

**Figure 2.**
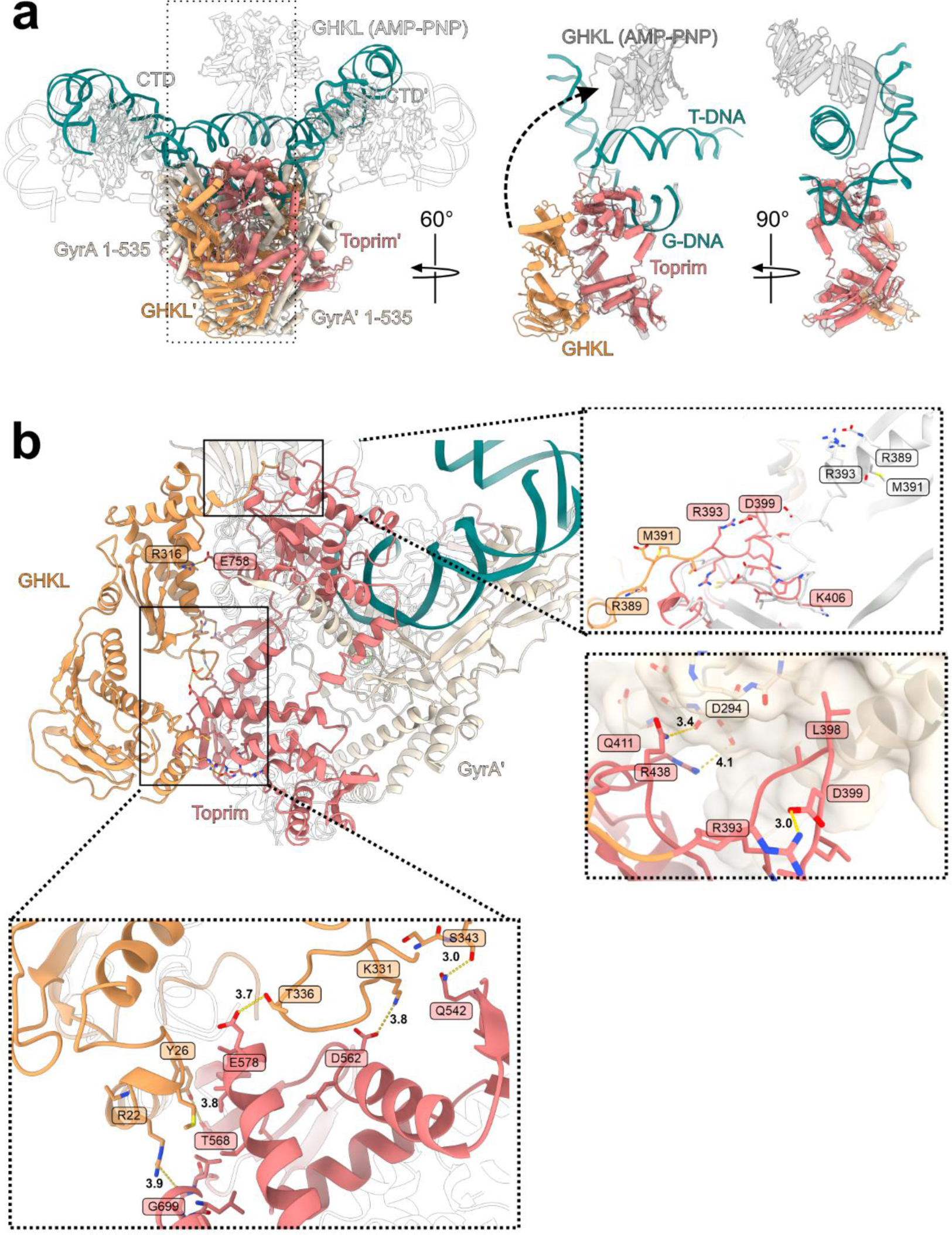
Position of the GHKL domain. **a,** Superposition of **Gyr-Mu217** (current work, color scheme as before) and E. coli gyrase in complex with 180 bp DNA, AMP-PNP and gepotidacin (PDB: 6RKW^6^, transparent contour). Boxed region (a single GyrB subunit) is shown in isolation in the right to illustrate the extreme motion of GHKL (12 nm shift & 180° rotation). AMP-PNP bound GHKL is shown in grey. **b,** An overall view of GHKL in downwards-folded conformation. Interactions with Toprim & loop conformation and interactions with GyrA Tower are shown as insets. AMP-PNP-bound structure (PDB: 6RKW) is shown as transparent contour or white cartoon (linker comparison between **Gyr-Mu217** and PDB:6RKW).

The extreme conformational change of GHKL is a result of the profound changes in the linker region (GyrB 386-406 in the current structure) (**Figure 2b**). The linker residues form a loop, stabilised by the salt bridge (R393-D399) and interactions with the Tower domain of the corresponding GyrA protomer. L398 of the linker occupies a hydrophobic pocket on GyrA surface while Q411 and R438 form hydrogen bonds to the Tower main chain (**Figure 2b**). In contrast, in the ADPNP-bound structure the linker is 10 residues long and extends in almost the opposite direction while residues 396-386 form a part of the extended transducer α-helix. This conformation is stabilised by multiple hydrogen bonds and salt bridges as shown in **Figure 2b**. The R393 residue is highly conserved, and amongst with the conserved lysines GyrA K284 and K308, and the residues from the blade 1 of the CTD forms a part of the ‘tunnel’ directing the T-segment along the top surface of the tetramer (see **Supplementary Figure 4d**). Therefore, GyrB R393 could be a sensor mechanism, coupling position of the GHKL with the T-DNA position.

Interestingly, a prior analysis by limited trypsin digestion suggested a special conformation stabilised by fluroquinolone (ciprofloxacin) binding that protects the GyrB 47 kDa domain (Toprim and insertion domain) from proteolysis^31^. Given that the protection has been observed only without ADPNP and lost upon ADPNP binding, we conclude that the protective conformation is likely resulting from GHKL domains folding down to protect a large surface area of GyrB47, as observed in our structure. We hypothesized that fluroquinolones may play a role in stabilisation of the chirally wrapped state, as binding of the drug would prevent strand passage. To investigate this, we have collected data on *E. coli* gyrase bound to the latest generation fluoroquinolone moxifloxacin (MFX; **Gyr-Mu217-MFX**). This resulted in a 2.8-Å structure displaying overall the same conformation as the drug-free complex (**Figure 3a; Supplementary Figures 5 and 6**) with the exception of the noteworthy changes required for cleavage of DNA and intercalation of the drug. This is to our knowledge the first structure of a full-length Gram-negative gyrase in complex with a fluoroquinolone. Both chains of DNA in the complex are cleaved (**Figure 3a**) to allow intercalation of two MFX molecules per gyrase complex in symmetry-related pockets (**Figure 3b**). A metal ion (interpreted as Mg^2+^ according to the buffer composition) connects the keto acid of the fluoroquinolone with S83 and D87 of GyrA subunit via a network of clearly visible water molecules resulting in the observed density for the Mg^2+^ ion having a characteristic octahedral shape (**Figure 3b,c**). Another contact is made by R121 from the catalytic dyad to the carboxyl of MFX (3.4 Å). The bicyclic C-7 substituent is protruding out from the DNA double helix to make contacts with E466 and K447 of GyrB: this explains previous biochemical data showing crosslinking of a chlorinated fluoroquinolone derivative to the E466C mutant^32^. The weaker interaction with S83 which is further than D87 from the water-metal ion bridge (3.6 Å vs 3.1 Å) is in line with the recent biochemical data showing that S83A mutation does not disrupt the MFX-induced DNA cleavage to the same extent as D87N does. S83, D87 and K447 are well-described as implicated in fluoroquinolone resistance^33^.

**Figure 3.**
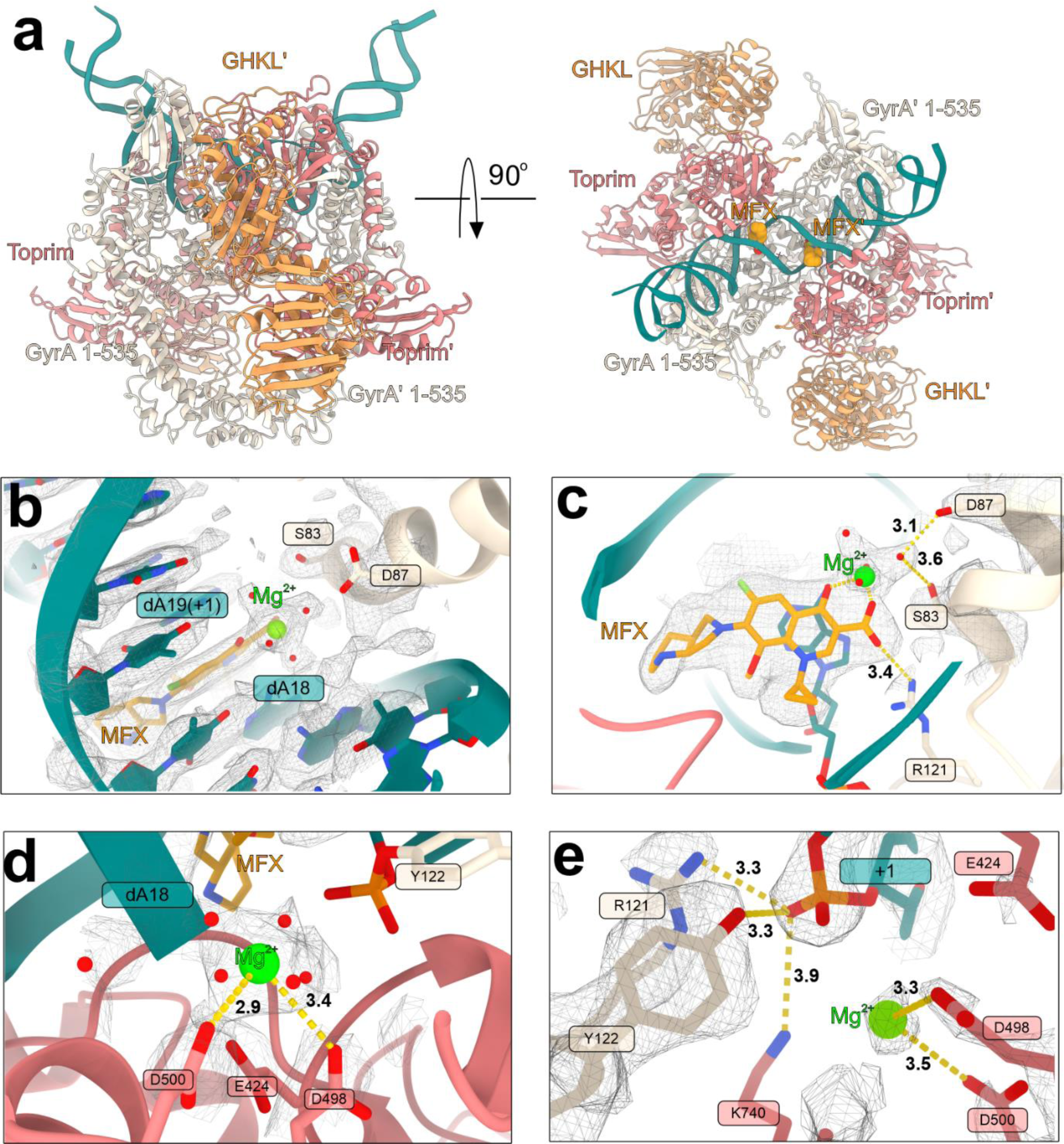
Structure of Gyr-Mu217-MFX and gyrase catalytic mechanism. **a**, A cartoon representation of the Gyr-Mu217-MFX atomic model (CTDs and chirally-wrapped DNA are not modelled). MFX is shown as golden spheres. **b**, Top view of MFX molecule binding pocket. MFX shown as golden sticks & magnesium ion as lime sphere. Density map contoured as 9σ. GyrA residues anchoring MFX molecule are indicated. **c,** Side view of MFX molecule; the water-metal ion bridge between MFX and S83/D87 of GyrA is shown. Distances in Å are indicated. Density map contoured at 9σ. **d**, catalytic metal ion position in **Gyr-Mu217** structure. Distances in Å towards closes residues are indicated. **e**, A catalytic site in one of the GyrA protomers (chain A) in **Gyr-Mu217**. Density is shown contoured at 15σ. Catalytic residues and corresponding distances are shown.

### Pre-cleavage gyrase complex contains a single metal ion

Comparison of the catalytic centres reveals interesting differences between the MFX-bound and drug-free structures. Surprisingly, there is almost no movement of GyrA protomers associated with DNA cleavage as for example was observed with the binding of a peptide-like drug albicidin. This observation helps to explain why fluoroquinolones are able to form complexes with heavily truncated enzymes (cleavage-reunion cores) and do not require DNA longer than 20 bp for stabilisation^34,35^. MFX binding and associated DNA cleavage requires a shift in the position of the nucleotide, accompanied by the formation of the phosphodiester bond between Y122 and DNA from both sides of the complex. A single metal ion is observed next to the catalytic tyrosine coordinated by D500 and D498 of GyrB but is situated too far to be competent for religation (so-called B-configuration^36,37^) (**Figure 3d & Supplementary Figure 7**). In the drug-free structure, a close comparison of the Coulomb potential density between Y122 and the scissile phosphate versus between the scissile phosphate and an adjacent nucleotide allows us to discern that the two DNA chains in **Gyr-Mu217** have subtly different conformations.

Both chains were modelled as uncleaved, but while a DNA strand next to the GyrA chain A (between dA18 and dA19;) could be modelled and refined well (**Figure 3e**), the Coulomb potential density for the scissile phosphate in the antiparallel chain (between dG21 and dT20 (5′-**T/G**ATTT-3’) cannot be refined with the uncleaved phosphate occupying the center of the observed density. The catalytic tyrosine is at 2.7 A distance from the phosphate in chain A, compared with the 3.3 A in chain C. Hence, we consider that chain C might represent an equilibrium between the pre-cleaved DNA and the initial state of cleavage with the phosphodiester bond formed between GyrA Y122 (chain C) and dG21. In the albicidin-stabilised Gyr-DNA complex^7^, the T/G pocket is larger and is the site of the drug intercalation hence the reason for the preferential cleavage might be the pre-existing stretch of DNA between these bases, as compared to the opposite strand. Given these subtle differences, we propose that the configuration we observe is very close to the actual pre-catalytic state. It involves stabilisation of the scissile phosphate by side-chains of GyrA R121 (3.3Å in chain A and 4.2 Å in chain C) and GyrB K740 (3.9Å) and interaction with the closely located single metal ion which we interpret as Mg^2+^(3.5Å). Interestingly, K740 density is less clear in the protomer where the DNA strand is intact. The K740A mutation was previously shown to be detrimental for enzyme activity and cause increased levels of cleavage^7^. Hence, we suggest that K740 is particularly important for DNA religation. Another observation in support of the pre-catalytic configuration comes from analysing the position of the catalytic metal coordinated by Toprim domain residues E424, D498 and D500. Two metals were previously simultaneously observed in a structure of a yeast type II topoisomerase, but all known structures of gyrase contained a single metal in one of the two configurations. Configuration A, where the metal primarily interacts with E424, is associated with intact DNA, and was observed for example in complexes with catalytic tyrosine mutated to phenylalanine, or in a complex of gyrase with the cleavage inhibitor LEI-800. Configuration B, where the metal interacts with D500, was observed in complexes with cleaved DNA, including this work (**Gyr-Mu217-MFX**). Surprisingly, the metal in **Gyr-Mu217** does not occupy either location and is most close to D498, a configuration closely resembling a drug-free cleaved structure of *S. pneumoniae* topoisomerase IV (**Supplementary Figure 7**). Observation of a single metal between the two previously observed configurations is compatible with the previously proposed mechanism where a single metal ion moves between three acidic GyrB residues^37^ while we cannot exclude temporary recruitment of a second ion to stabilise the catalytic intermediate complex. Upon DNA cleavage, the small movements associated with the nucleotide shift, K740 and metal repositioning will result in small shifts of corresponding GyrB TOPRIM loops (498-502) and (738-742); these small changes are propagated by the TOPRIM and particularly by the TOPRIM insert resulting in 6-Å distances between the α-helices in this region (between the **Gyr-Mu217** and **Gyr-Mu217-MFX**), enabling transmission of the DNA cleavage status to the GHKL domains and other parts of the enzyme.

### Ratchet and pawl: a mechanistic model for gyrase motor

Two main frameworks have been traditionally used to explain how molecular motors operate: the “power stroke” and the “Brownian ratchet”^1^. In the power stroke models, ATP consumption generates a large free energy gradient across the motor step distance, resulting in an irreversible transition. The Brownian ratchet is driven by thermal fluctuations making the motor visit previous and forward states, while conformational changes triggered by ATP binding and hydrolysis increase the affinity toward forward states, resulting in an overall directionality. According to the node inversion mechanism proposed more than 40 years ago^9,10^, the directionality of gyrase results from the chiral selection, and the input of energy of ATP that is converted into mechanical energy used to drive the unidirectional strand passage. In this work, we have determined the molecular mechanism which is responsible for this chiral selection and showed that without energetic constraints, gyrase indeed stabilises a positively supercoiled DNA loop. We propose the following model for the gyrase motor which is illustrated in **Figure 4**:

**Figure 4.**
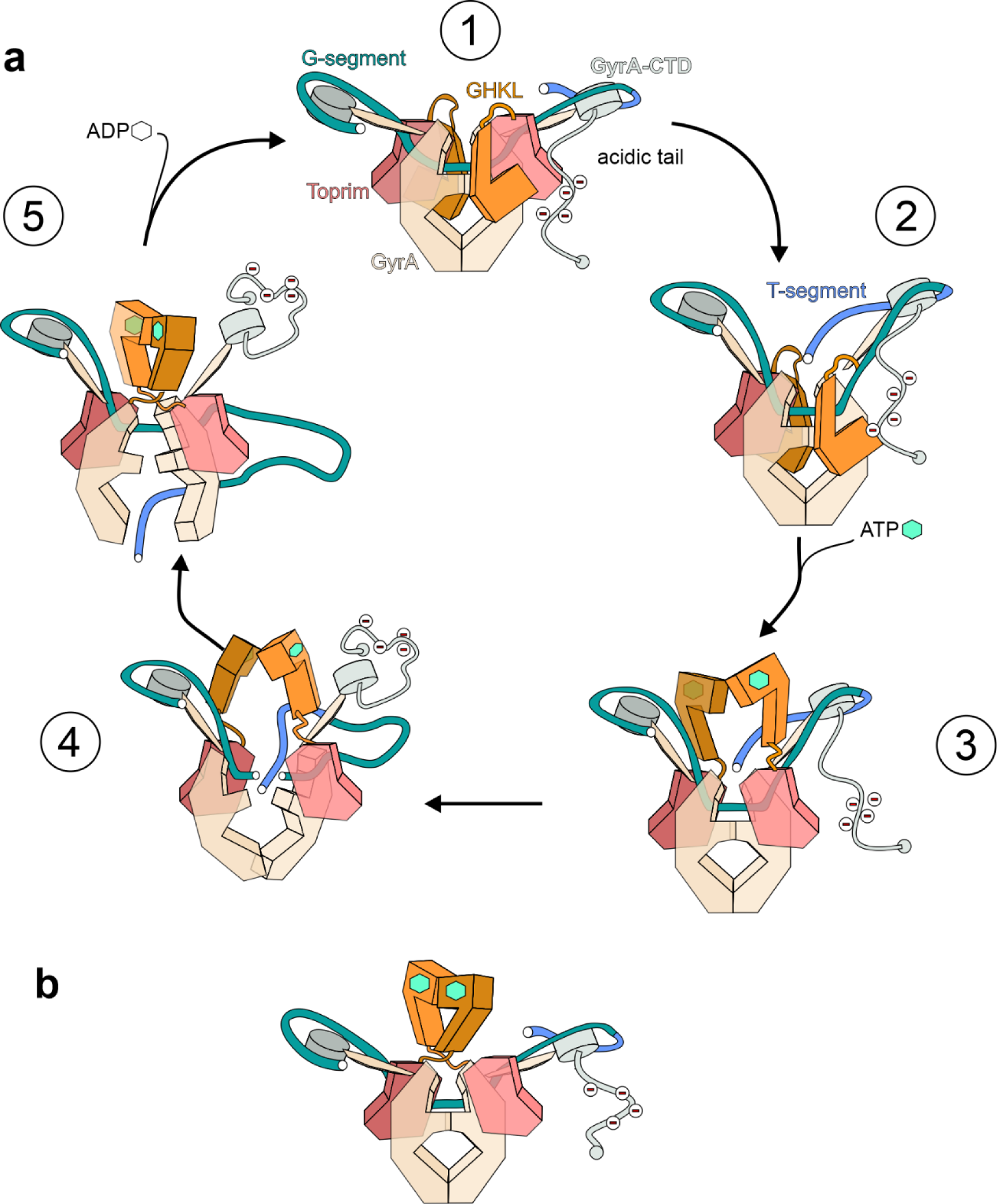
A ratchet & pawl model for gyrase strand passage mechanism. **a,** The catalytic cycle consists of 5 stages as described in the paper. Nucleotide binding is indicated by a green hexagon (ATP) or empty hexagon (ADP); acidic tail of CTD is indicated by negative charges. Stage 1 – Ω-state occurring after initial DNA binding or immediately after enzyme reset. Stage 2 - α-state where DNA is constrained in a (+) supercoiled loop and the catalytic center is primed for cleavage as in **Gyr-Mu217** (current paper). Stage 3 – following ATP binding, the GHKL domains disengage and move upwards to undergo Brownian movement towards the lowest energy conformation. They might be guided towards T-DNA by charge attraction. At the same time, GHKL movement might release the acidic tail, resulting in the loss of wrap. Stage 4 – GHKLs follow the moving T-segment in its thermal excursion downwards and prevent reversal of strand passage as it occurs. The probability of the event depends on the potential energy of DNA. Stage 5 – after completion of strand passage, the T-segment can only escape through the bottom gate, resulting in an overall change of linking number by −2 and completion of the cycle. Nucleotide hydrolysis in the absence of the chiral wrap allows enzyme reset. In case of antibiotic rescue factors QnrB1 and MfpA, this reverse motion of GHKLs might power the release of an antibiotic from the complex. **b,** a trapped post-strand-passage state with dimerised GHKL module as seen in the AMP-PNP bound cryoEM structures occurs when the enzyme reset cannot happen.

Initial binding of gyrase to DNA might result in an Ω state intermediate, which is not chirally wrapped (**1**) and was visualised previously^6^. We propose that our chirally wrapped structure represents the initial stage of the reaction, or a pre-strand passage α intermediate (**2**). It has been shown that chiral wrap occurs with high efficiency on all substrates (linear, negatively and positively supercoiled DNA) and is enabled by tight binding of DNA to β-pinwheels; the wrapped state has been also shown to be the most populated state in rotor bead tracking experiments^19^. A Mu217 sequence might be particularly conducive for the wrapping, facilitating α state formation. To enable progressive negative supercoiling on substrate of increasing superhelicity, DNA binding to CTDs is very tight; it has been shown that an artificial increase in CTD positive charges stabilises the wrap, allowing a small increase in maximal superhelicity obtained but a large decrease in supercoiling speed^38^. Thus, as was originally proposed in 1979-1980, it is the transition to the next stage that ultimately requires energy input in form of the energy of ATP. After binding of ATP, conformational changes around the active site (previously described in the literature rearrangement of the ‘switch loop’, **Supplementary Figure 8**) result in an 11° rotation of transducer domain^39–43^, that trigger disengagement and undocking of GHKLs. In this scenario, ATP binding energy compensates for the loss of bonds between the GHKL and Toprim insert. While it could be conceived that elastic energy stored in the “folded” GHKL enables the 12 nm shift and mechanical push of DNA downwards, we find this extremely unlikely. Not only does the compact GHKL not have any obvious reservoir for such mechanical energy, but it would also be quickly dissipated in the conditions of high drag and thermal collisions experienced by the motor. In our model, the energy of ATP binding enables a large initial conformational transition followed by thermal relaxation to the next lower potential well where GHKL is rotated 180° as observed in the AMP-PNP-bound structure (**3**). This relaxation requires the T-segment to move beyond the plane of the G-segment, which in turn requires DNA cleavage. It is shown that gyrase naturally maintains an equilibrium between cleaved and intact DNA which is normally shifted towards relegation; however, nucleotide binding stimulates DNA cleavage by a yet undetermined mechanism. The T-segment passage requires a removal of the DNA wrap, and indeed that what happens upon the nucleotide binding^44^. The mechanism for this loss of wrap might include a conserved acidic tail which was shown to be essential for *E. coli* gyrase supercoiling. While the tail is unstructured, and not observed in our maps, we propose that it initial position (perhaps interacting with docked GHKLs) allows CTDs to move upwards and fully engage DNA, stabilising a positive supercoil. The events associated with the nucleotide binding might simultaneously disengage the acidic tail, which in turn facilitates removal of wrapped DNA from the CTD. Thus, effective supercoiling would require tight coordination of nucleotide binding with both loss of wrap and DNA cleavage to create a brief window of opportunity, during which T-DNA strand passage (**4**) can happen. While the molecular basis of this coordination is not determined, it might be controlled by GHKL position.

It is well established that the probability of strand passage per round of nucleotide binding depends on the nature of the substrate (almost 1 for positively supercoiled DNA and 0 for negatively supercoiled DNA). Therefore, in our model strand passage is a random event, the probability of this event depending on temperature and the DNA torsional energy. After a successful strand passage attempt, the T-DNA hovers below the DNA-gate plane, allowing the GHKL to fully rotate (**5**) resulting in the Ω conformation with the ATPase domains dimerising. Rotation and dimerization prevents the reversal of the process (upwards escape of the T-DNA) ultimately ensuring reaction directionality. With AMP-PNP, the dimer remains irreversibly locked and is observed as such by cryoEM (**Figure 4b**), while in the course of a normal reaction, dimerization induces ATP hydrolysis that in our model relaxes GHKL to their initial ‘folded’ conformation. Slow hydrolysis of ATP by a monomeric GHKL ensures that the enzyme has multiple attempts for strand passage even on negatively supercoiled substrates.

Remarkably, it has been shown that an enzyme that is incapable of cleaving DNA does not demonstrate DNA-stimulated increase in the rate of ATPase activity. It is also noteworthy that binding of the peptide inhibitors albicidin or microcin B17 requires a strand passage attempt and the binding (but not hydrolysis) of the nucleotide; at the same time, hydrolysis of nucleotide was shown to be important for activity of antibiotic resistance factors QnrB1 and MfpA^29,30^. While the peptide toxins could occupy the larger space between dissociated GyrA protomers occurring after strand passage, they are probably using the motion of relaxing GHKLs to remove bound drugs (**Supplementary Figure 3**).

A crucial difference between our model and previously proposed schemes (e.g.^45–47^) is the observation that the GHKL dimerization simply cannot happen prior to strand passage, with the traditional model of a clamp-like T-segment capture sterically incompatible with the observed position of T-DNA. We propose that rather than actively pushing or capturing the T-DNA, GHKL rotation acts as an irreversible conformation change (a ‘pawl’), ensuring the directionality of the probability-based Brownian ratchet mechanism. Therefore, in its activity *E. coli* gyrase combines the elements of power stroke (a large movement connected with the ATP binding) with the ratchet-and-pawl like mechanism (a strand passage event which is made irreversible by a conformational change).

Our model allows us to make important predictions regarding the sequence of events and role of individual gyrase subunits and interfaces. Particularly interesting would be the application of non-interfering, in solution techniques such as smFRET^48^ or EPR to directly observe predicted conformational changes. These experiments could also investigate the proposed key role of the C-tail in movement between different conformational stages. Pioneering work on construction and testing of heterocomplexes having domains inactivated on a single side has already resulted in important observations^16,49^. Supercoiling set points are different even in closely related organisms, and likely even more different in thermophilic bacteria and archaea^50,51^. We believe that the structural and theoretical framework proposed in this manuscript will spring further fruitful discussions towards fundamental understanding and practical use of gyrase and other molecular motors.

## Methods

### CryoEM sample preparation

*E. coli* GyrA and GyrB proteins were purified as previously described^7^ using metal affinity, Strep-tag and ion-exchange chromatography. Proteins were concentrated to 12 mg/ml prior to complex formation and dialysed overnight into cryo-EM buffer [25 mM Na-HEPES pH 8, 30 mM potassium acetate, 2.5 mM magnesium acetate, 0.5 mM Tris (2-carboxyethyl) phosphine (TCEP)] in presence of equal molar amount of Mu217 DNA in a Pur-A-Lyzer Mini (Merck). Mu217 DNA was purified as previously described^7^. For MFX complex, drug was added to the dialysis buffer at 50 µM concentration. Dialysed sample was concentrated to 15 µM, additionally supplemented with 100 µM moxifloxacin and incubated at 37 °C for 15 minutes. Before grid freezing, CHAPSO (8 mM) was added and the samples were spun at 21 000g for 60 min. 4 µl of sample was applied to the Quantifoil (R2/1, 300 copper mesh) glow-discharged grids. Grids were blotted for 6 s and plunge-frozen in liquid ethane using Vitrobot Mark IV (at 95% humidity, 10 °C).

### CryoEM data collection & analysis

CryoEM data were collected on Krios G3i microscope at Polish national cryoEM facility SOLARIS using Gatan K3 camera with Gatan BioQuantum energy filter operated with a slit width of 20 eV. Movies were collected at a 105 000× nominal magnification, resulting in a calibrated physical pixel size of 0.86 Å using EPU v2.10.0.1941REL. Movies were saved at physical pixel size as gain-corrected TIFF files. For **Gyr-Mu217**, 8 508 movies were collected with the range of defoci set as −1.8, −1.5, −1.2, −0.9 μm and a total dose of 40.22 e/Å^2^ over 40 frames. 8 405 movies were kept for further processing in CryoSPARC v. 4.2.1^52^. Movies were motion and CTF corrected in patch mode. 658 859 particles were picked using cryoSPARC template picker and extracted with a pixel size of 1.72 Å/px. Binned particles underwent 2 rounds of 2D classification to yield cleaned stack of 230 444 particles. Ab initio job was used to classify in 3D, followed by a non-uniform refinement^53^. Particles were re-extracted at physical pixel size and refined correcting for local defocus yielding a 2. Å consensus map. Particles underwent a round of reference-based motion correction^54^ as implemented in cryoSPARC, followed by heterogenous refinement with two classes (a map and a low-passed filtered map) to remove particles that did not contribute to high resolution structure. After a second round of polishing, 3^rd^ and 4^th^ order CTF aberrations correction^55^ and Ewald sphere correction^56^, the final resolution was 2.32 Å after non-uniform refinement in cryoSPARC. To further improve density for the CTD in the map, 3D classification without alignment was carried out with 10 classes, using mask around a CTD. Local refinement of particles from 3 best classes yielded a 2.94 Å map which was combined with the consensus map using ChimeraX *vop maximum* command for a composite map used for refinement.

For **Gyr-Mu217-MFX**, 4 500 movies were collected using the range of defoci set as −2.1, −1.8, −1.5, −1.2, −0.9 μm and a total dose of 40.68 e/Å^2^ over 40 frames. 4 246 were kept for further processing. 190 069 particles were picked using Topaz^57^ and extracted with a pixel size of 1.72 Å/px. Binned particles underwent a round of 2D classification yielding 152 001 particles, and a round of 3D classification (Ab initio) yielding 133 625 particles. After re-extraction, refinement and a reference-based local motion correction as implemented in cryoSPARC 4.4, followed by a non-uniform refinement with correcting for local defocus, 3rd and 4th order CTF aberrations and Ewald sphere^55^, a consensus map was obtained with a resolution of 2.46 Å used for the refinement of the core enzyme (GyrA 7-524; GyrB 405-804). This map displayed heterogeneity in the position of GyrA CTDs and GyrB GHKL domains; to address that, a mask was applied around GHKL domains (**Supplementary Figure 5**) followed by classification without alignment in cryoSPARC (5 classes). Classes with GHKL density predominantly from one or another side of the core complex were obtained; symmetrical classes were combined together and refined to yield a 2.61 Å map used for building of the model that incorporated GyrB GHKL and GyrA CTD domain.

### Model building and refinement

Model for cleavage-reunion core was manually built in Coot^58^ guided by a map processed by deepEMhancer^59^ and based on the previously available high-resolution structures (PDB: 7Z9C ^7^). The GHKL domain was manually built based on available crystal structure (PDB:1EI1^39^). The C-terminal domain was built using crystal structure PDB: 1ZI0^23^ and ModelAngelo^60^ followed by manual geometry optimisation in Coot. Poor resolution regions were refined using ISOLDE^61^. To build DNA, bases around the cleavage site were manually assigned and the rest of the wrapped DNA was constructed using ideal B-form DNA blocks in Coot. cryoREAD-generated model ^62^ was used for guidance and to verify DNA positioning. Complete model was refined in real space using Phenix^63^ against an unsharpened map with secondary structure restraints for protein and DNA bases. NCS restraints were used during first few rounds of refinement for stabilisation and subsequently switched off. All visualisation, superposition & surface calculation was done in ChimeraX^64^. To build the **Gyr-MFX-Mu217** (**2**), GHKL domain and CTD were copied from the **Gyr-Mu217**, rigid-body fitted and refined in real space.

## Data availability

All data needed to evaluate the conclusions in the paper are present in the paper or available from the authors upon reasonable request. The **Gyr-Mu217** and **Gyr-Mu217-MFX** coordinates have been submitted to the Protein Data Bank (https://www.rcsb.org/) with PDB IDs XXX, respectively. Corresponding EM maps have been submitted to the Electron Microscopy Data Bank (https://www.ebi.ac.uk/pdbe/emdb/) with IDs EMD-XXX, respectively.

## Acknowledgments

We thank the national cryoEM facility and staff at the SOLARIS National Synchrotron Radiation Centre, Poland (Polish Ministry of Education and Science project: ‘Support for research and development with the use of research infrastructure of the National Synchrotron Radiation Centre SOLARIS’ under contract number 1/SOL/2021/2). We thank M. Rawski, P. Indyka & Grzegosz Ważny for the excellent technical support. We acknowledge MCB Structural Biology Core Facility (supported by the TEAM TECH CORE FACILITY/2017-4/6 grant from the Foundation for Polish Science). E.M., M.P. and J.G.H. acknowledge the funding from National Science Centre (NCN), Poland – OPUS20: 2020/39/B/NZ1/02898. D.G. is a Wellcome Trust Sir Henry Dale Fellow (221868/Z/20/Z); work in his lab is also supported by the BBSRC funded Institute Strategic Programme “Harnessing Biosynthesis for Sustainable Food and Health” (HBio) (grant number BB/X01097X/1). We thank Anthony Maxwell & James Berger for feedback on the manuscript before publication.

## SUPPLEMENTARY INFORMATION

**Supplementary Table 1.** CryoEM data collection & refinement statistics.

|  | EcGyr-Mu217 | EcGyr-Mu217-MFX |  |
| --- | --- | --- | --- |
| **Data collection and processing** |  |  |  |
| Microscope | ThermoFisher Krios G3i | ThermoFisher Krios G3i |  |
| Magnification | 120,000× | 120,000× |  |
| Voltage (kV) | 300 | 300 |  |
| Electron dose (e-/Å<sup>2</sup>) | 40.22 | 40.68 |  |
| Detector | Gatan K3 | Gatan K3 |  |
| Defocus range (-μm) | 1.8-0.9 | 2.1-0.9 |  |
| Pixel size (Å) | 0.86 | 0.86 |  |
| Symmetry imposed | C1 | C1 |  |
| Micrographs (no.) | 8 508 | 4 500 |  |
| Initial particle images (no.) | 658 859 | 190 069 |  |
| Final particle images (no.) | 170 369 | 132 931<br>(consensus map) | 79 415<br>(GHKL+CTD) |
| Global map resolution (Å) | 2.3 | 2.5 | 2.6 |
| FSC threshold | 0.143 | 0.143 |  |
| **Refinement** |  |  |  |
| Model resolution (Å) | 2.57 | 2.6 (core) | 2.9<br>(core+GHKL+CTD) |
| FSC threshold |  | 0.5 |  |
| Map sharpening *B* factor (Å<sup>2</sup>) | No sharpening performed | -55.5 (core) |  |
| *Model composition* |  |  |  |
| Non-hydrogen atoms | 31791 | 16420 | 24770 |
| Protein residues | 3237 | 1836 | 2915 |
| Nucleotides | 312 | 84 | 84 |
| Ligands | 2 | 6 | 6 |
| *Mean B factors (Å<sup>2</sup>)* |  |  |  |
| Protein | 149.87 | 69.07 | 138.28 |
| Nucleotide | 202.18 | 40.84 | 71.79 |
| Ligands | 91.72 | 36.88 | 71.35 |
| *R.m.s. deviations* |  |  |  |
| Bond lengths (Å) | 0.002 | 0.003 | 0.004 |
| Bond angles (°) | 0.457 | 0.509 | 0.525 |
| **Validation** |  |  |  |
| MolProbity score | 1.43 | 1.33 | 1.85 |
| Clashscore | 4.25 | 2.91 | 4.82 |
| *Ramachandran plot* Favored (%) | 96.59 | 96.76 | 95.27 |
| Allowed (%) | 3.41 | 3.24 | 4.73 |
| Disallowed (%) | 0.00 | 0.00 | 0.00 |

**Supplementary Figure 1.**
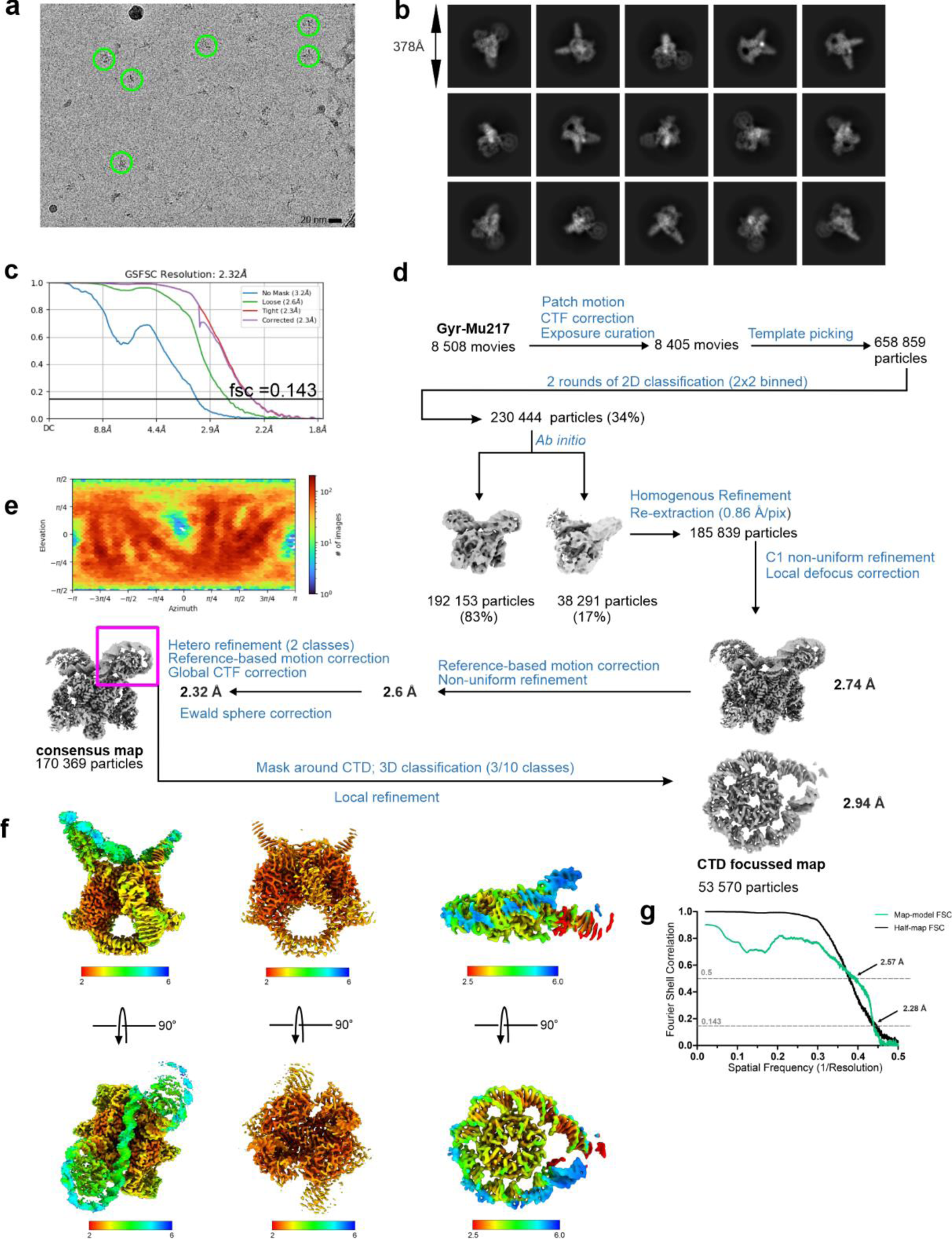
CryoEM data processing scheme for Gyr-Mu217. **a**, A representative motion-corrected micrograph, gyrase particles are encircled. **b**, A selection of 2D classes, box size in angstroms indicated. **c,** FSC curve for the final reconstruction as output by cryoSPARC. **d**, Processing scheme (see Methods for description). **e**, Euler angle distribution as output by cryoSPARC. **f**, Local resolution maps illustrate resolution distribution from 2.1 Å next to the DNA, to >5 Å towards the ends of the wrapped DNA molecule & flexible CTD domains. **g**, Map-to-model FSC curve (green) compared with half-maps FSC curve (black) as output by Phenix (soft mask based on the atomic model is used). FSC=0.5 (for map-to-model FSC) and FSC=0.143 (for half-map FSC) values are indicated with arrows.

**Supplementary Figure 2.**
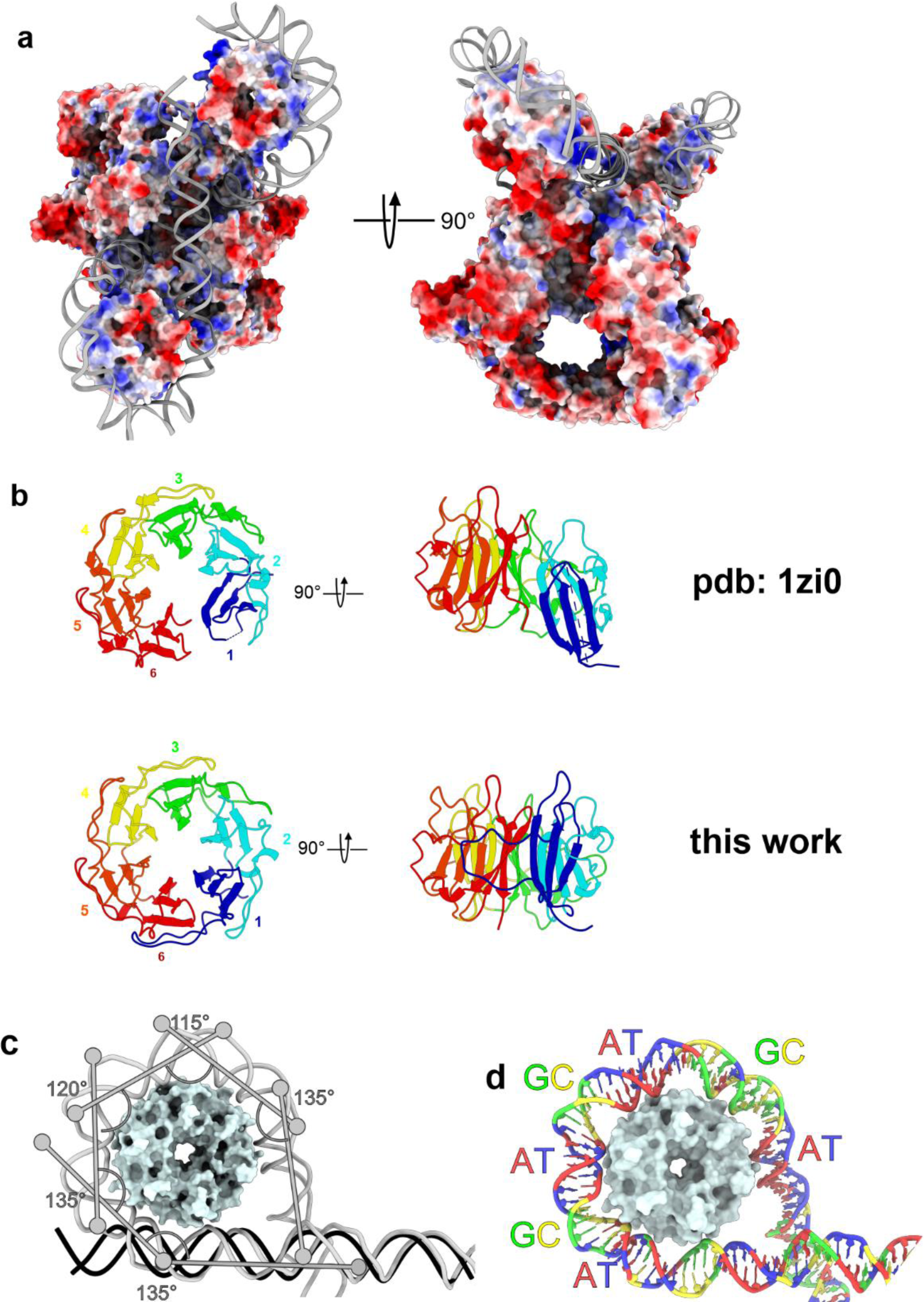
Gyr-Mu217 surface charge. **a,** A molecular surface representation of Gyr-Mu217 model colored by Coulombic potential as calculated by ChimeraX module using default parameters. DNA is shown as cartoon grey representation. Blue corresponds to positive charge and red to negative charge. A positively charged guiding pathway for the T-segment is visible. **b**, a comparison of b-pinwheel domains modelled in this work (Gyr-Mu217) and based on the structure of an isolated CTD ^23^(PDB: 1ZI0). Individual blades are colored. **c**, an illustration of DNA bending around the CTD. 5 bends introduce a total ∼260° angle in DNA. **d**, Sequence-specificity of DNA wrapping. AT-rich parts of the Mu217 right arm are positioned to form the minor groove facing the protein, while the GC-rich minor grooves face outwards, reminiscent of a nucleosome DNA wrap.

**Supplementary Figure 3.**
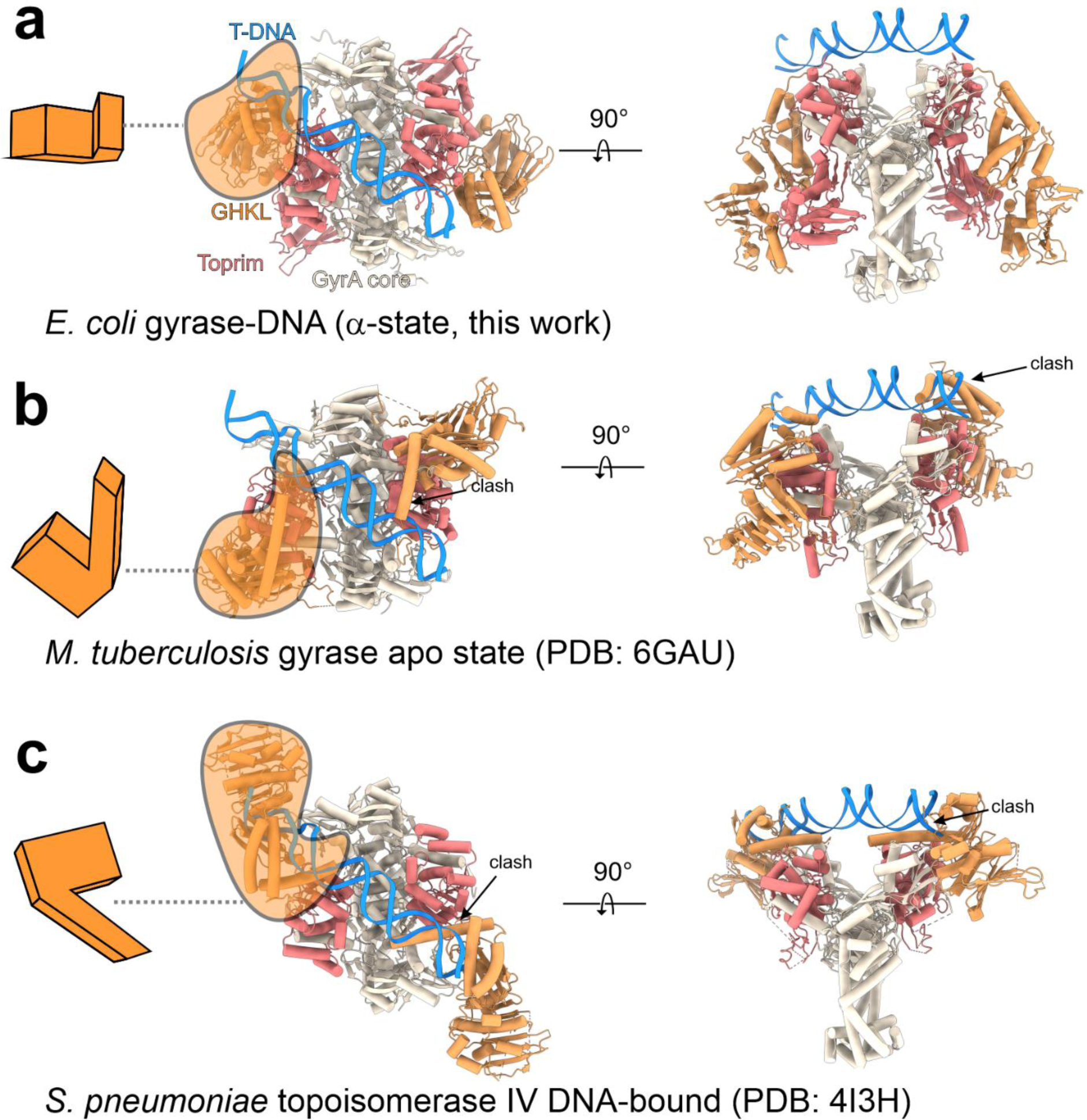
Comparison of the position of GHKL domains in different type IIα topoisomerase structures. **a, Gyr-Mu217** (current work). An orange cartoon at the left illustrates the orientation of the GHKL: the ‘inner’ surface faces the reader. The same coloring scheme is used as in other main figures. **b**, an ‘open-clamp’ *M. tuberculosis* gyrase structure (PDB:6GAU) is superimposed based on the GyrA protein, whilst the T-segment orientation is kept intact. GHKL is facing the ‘side’ surface to the reader, and clashes with the T-segment. **c**, an *S. pneumoniae* topoisomerase IV (PDB:4I3H) is superimposed based on the ParC protein, while the position of the T-segment is kept. Once again, the GHKL is clashing with the T-segment.

**Supplementary Figure 4.**
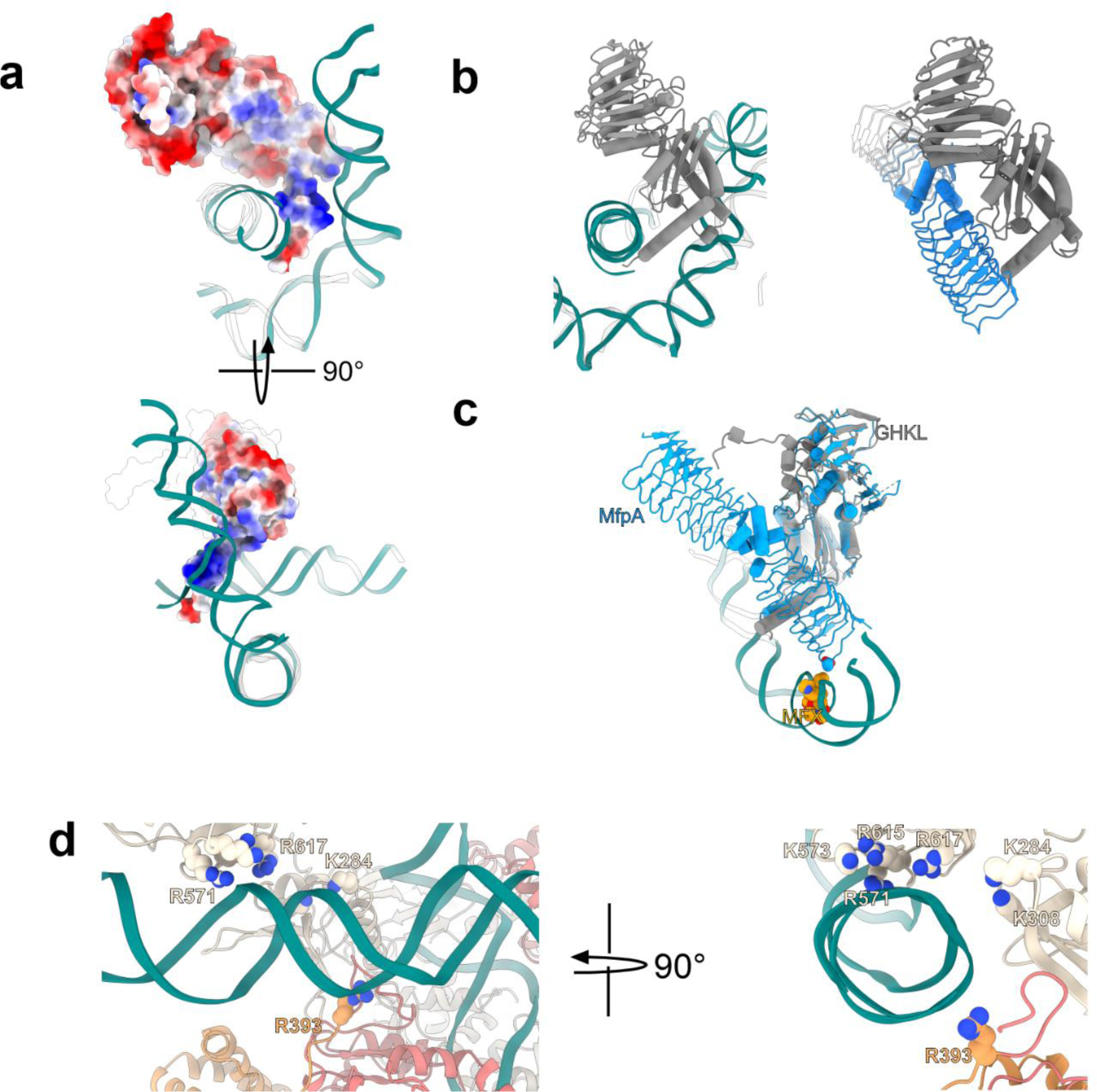
Comparison of the nucleotide-bound GHKL domain position relative to the T-DNA and T-DNA mimic, antibiotic resistance protein MfpA. **a,** A superposition of GHKL bound to AMP-PNP (only one monomer is shown for clarity, PDB:6RKW) and positively supercoiled DNA loop from Gyr-Mu217 (current study, colored teal). GHKL is shown as Coulomb potential colored surface representation. Two DNA binding surfaces are visible. **b**, (*left*) A superposition of GHKL bound to AMP-PNP (PDB: 6RKW, GHKL is gray and DNA transparent) and positively supercoiled DNA loop from **Gyr-Mu217** (current study, colored teal) and a comparison with (*right*) *Mycobacterium smegmatis* GHKL in complex with MfpA in the same orientation (PDB:6ZT5). Note the similarity in position of MfpA molecule and T-DNA. **c**, A superposition of MfpA-GyrB47 crystal structure (PDB: 6ZT5, blue) and **Gyr-Mu217-MFX** (current work). MFX and MfpA Asp24 are shown as van der Waals spheres. MfpA can reach MFX molecule, offering a potential explanation for the nucleotide hydrolysis-dependent resistance mechanism. **d**, an illustration of the role of GyrA Tower and GyrB linker domains in forming a ‘tunnel’ for the T-segment.

**Supplementary Figure 5.**
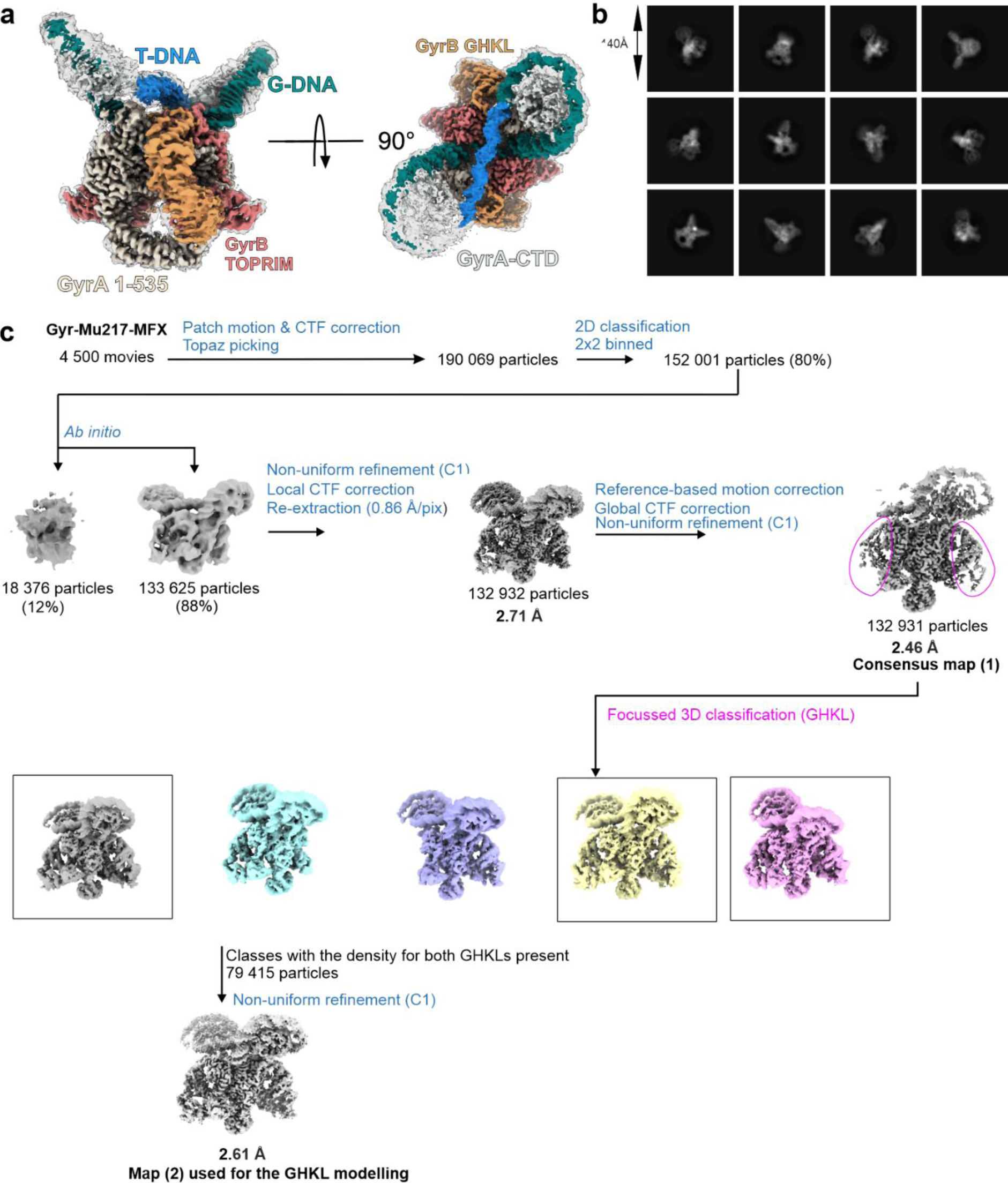
CryoEM data processing scheme for Gyr-Mu217-MFX. **a**, Different views of the cryoEM density map for the **Gyr-Mu217-MFX** complex presented at two contour levels (9σ and 5σ). The 9σ map is colored according to the color scheme used elsewhere in the manuscript. **b**, A selection of 2D classes, box size in angstroms indicated. **c,** Processing scheme (see *Methods* for description).

**Supplementary Figure 6.**
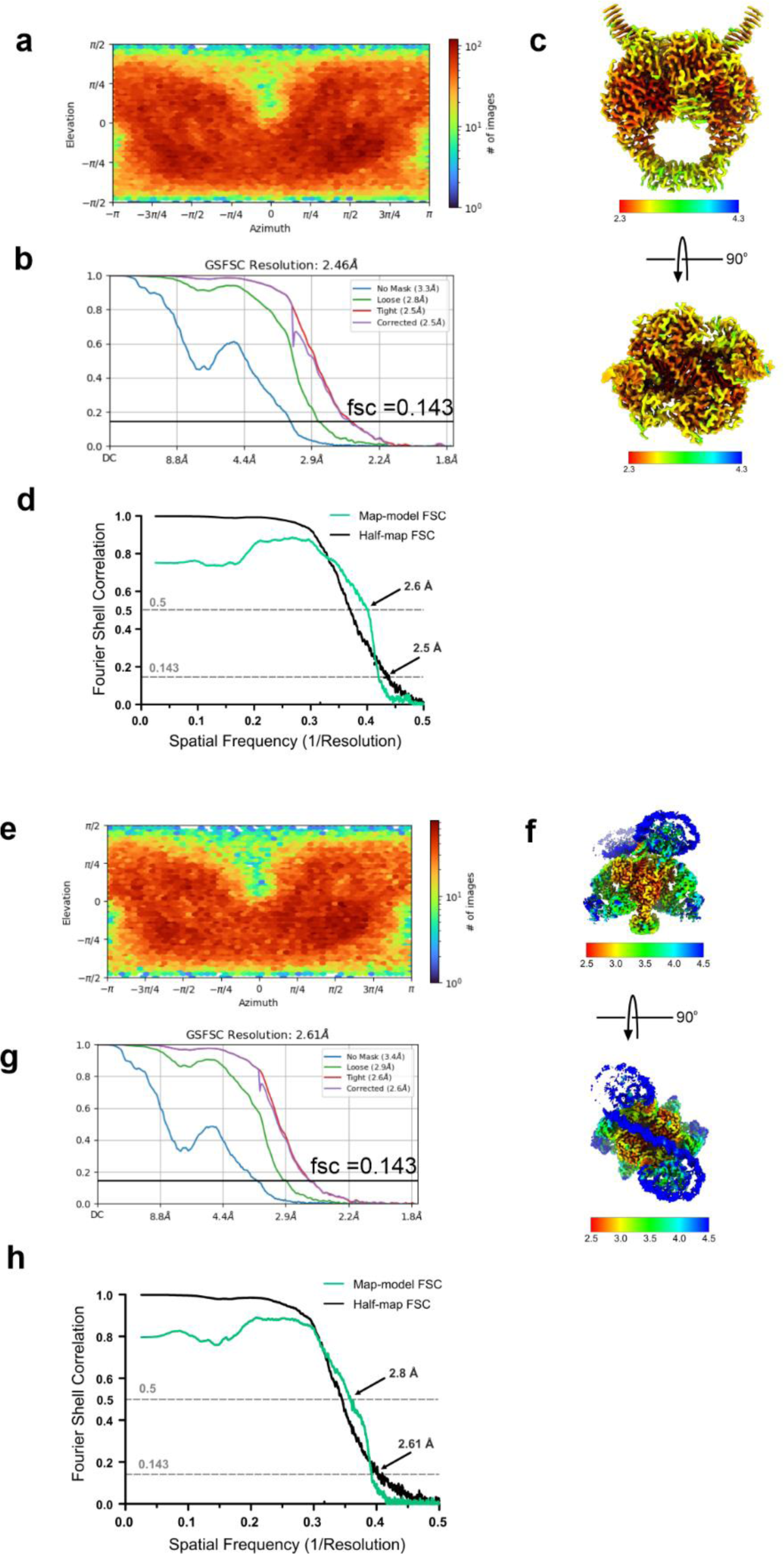
CryoEM map validation for Gyr-Mu217-MFX. **a**, Euler angle distribution as output by cryoSPARC. **b**, FSC curve for the consensus reconstruction as output by cryoSPARC. **c**, Local resolution map for the consensus reconstruction (contoured ∼11σ). **d**, Map-to-model fit curve. **e**, Euler angle distribution as output by cryoSPARC. **g**, FSC curve for the focussed reconstruction as output by cryoSPARC. **f**, Local resolution map for the consensus reconstruction (contoured ∼9σ). **h**, Map-to model fit

**Supplementary Figure 7.**
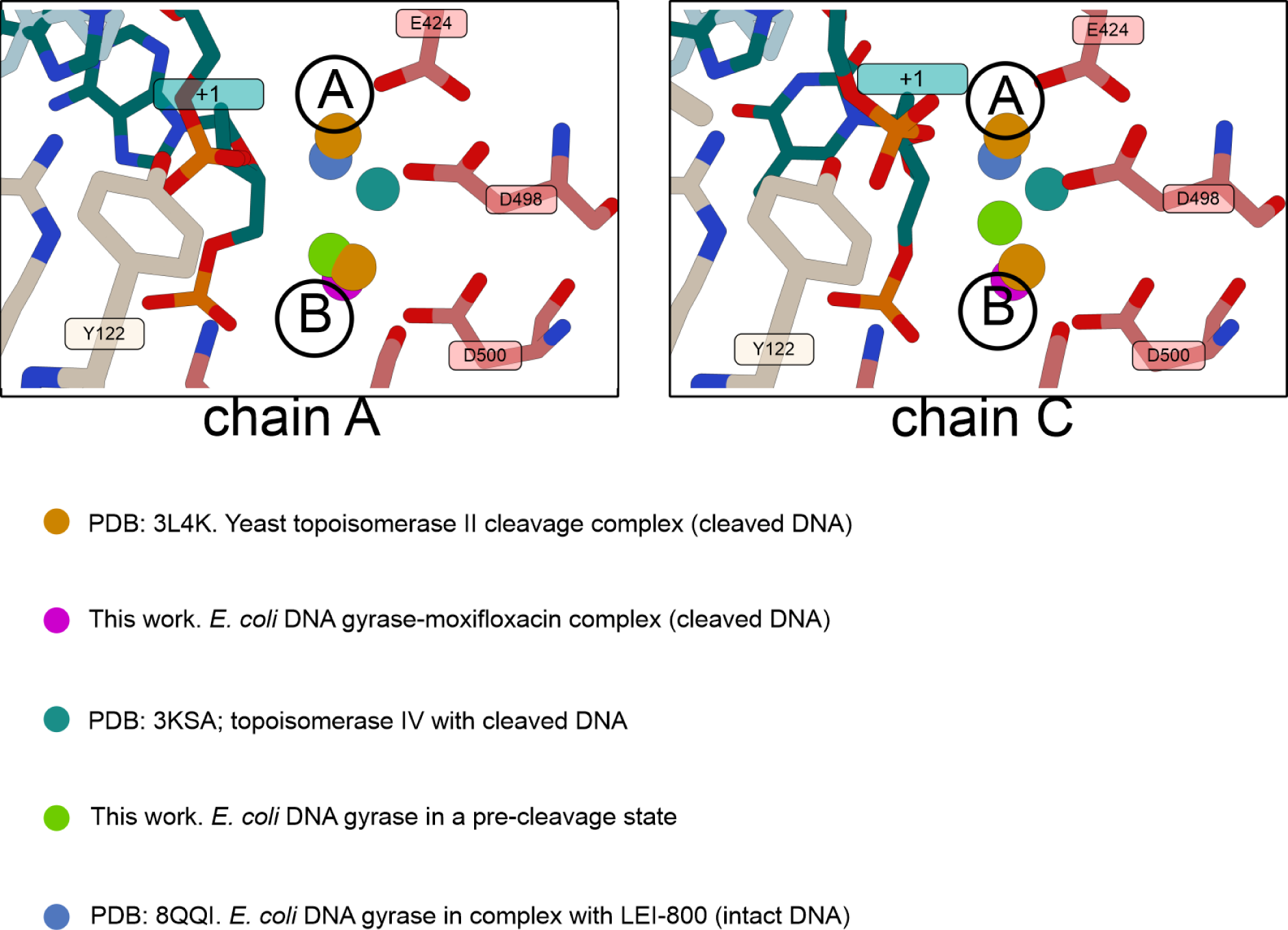
Comparison of metal ion position in Gyr-Mu217 and other type II topoisomerase structures. Catalytic tyrosine, scissile phosphate and Toprim catalytic triad are shown as sticks. Chain A and Chain C in **Gyr-Mu217** are modelled separately due to the subtle differences between the chains. All other structures are superimposed based on Toprim domain using ChimeraX matchmaker tool. A and B -type coordination is shown: in structures with uncleaved DNA, metal is clustered in the A site while in structures with cleaved DNA, in B site. **Gyr-Mu217** places metal closer to the middle position which is similar to what was reported for the drug-free structure of topoisomerase IV with cleaved and resealed DNA.

**Supplementary Figure 8.**
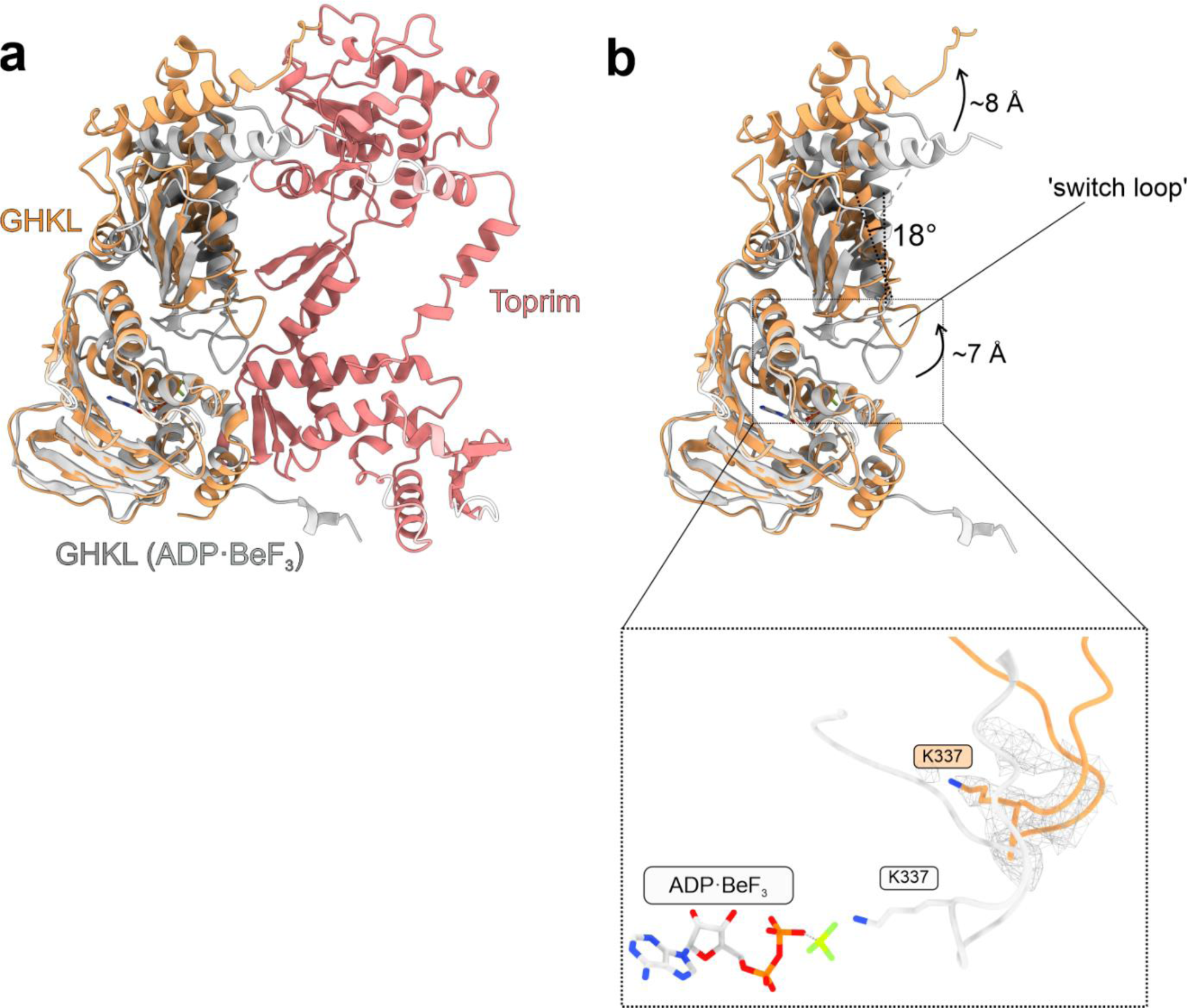
‘Switch loop’ conformation in nucleotide-free structures. **a.** A superposition of the ATPase domain (GHKL-transducer) in ADP:BeF_3_ structure of an isolated domain (PDB: 4U9) (white cartoon representation) and the conformation of the same domain in the chirally wrapped (α-state) holocomplex (**Gyr-DNA**, this study). GyrA subunits, opposite GyrB subunit and DNA not shown. **b**. Same as **a**, but Toprim removed for clarity, while movements of the transducer helices highlighted by arrows. Inset shows the ‘switch loop’ in the isolation. in the ADP:BeF_3_ structure (PDB: 4U9). K337 is interacting with BeF_3_. Orange cartoon representation shows the equivalent ‘switch loop’ in the **Gyr-DNA** structure (this work). Density (gray mesh) around the loop is shown, supporting modelled lysine orientation.

